# LIFE-Seq: A universal Large Integrated DNA Fragment Enrichment Sequencing strategy for transgene integration in genetically modified organisms

**DOI:** 10.1101/2021.09.07.459346

**Authors:** Hanwen Zhang, Rong Li, Yongkun Guo, Yuchen Zhang, Dabing Zhang, Litao Yang

## Abstract

Molecular characterisation of genetically modified organisms (GMOs) yields basic information on exogenous DNA integration, including integration sites, entire inserted sequences and structures, flanking sequences and copy number, providing key data for biosafety assessment. However, there are few effective methods for deciphering transgene integration, especially for large DNA fragment integration with complex rearrangement, inversion, and tandem repeats. Herein, we developed a universal <u>L</u>arge Integrated DNA <u>F</u>ragments <u>E</u>nrichment strategy combined with PacBio <u>Seq</u>uencing (LIFE-Seq) for deciphering transgene integration in GMOs. Universal tilling DNA probes targeting transgenic elements and exogenous genes facilitate specific enrichment of large inserted DNA fragments associated with transgenes from plant genomes, followed by PacBio sequencing. LIFE-Seq were evaluated using six GM events and four crop species. Target DNA fragments averaging ∼6275 bp were enriched and sequenced, generating ∼26,352 high fidelity reads for each sample. Transgene integration structures were determined with high repeatability and sensitivity. Compared with whole-genome sequencing, LIFE-Seq achieved better data integrity and accuracy, greater universality, and lower cost, especially for transgenic crops with complex inserted DNA structures. LIFE-Seq could be applied in molecular characterisation of transgenic crops and animals, and complex DNA structure analysis in genetics research.

## Introduction

In recent decades, many genetically modified (GM) crops have been developed and approved for plants and plant products, resulting in economic and environmental benefits from higher yields, more efficient use of natural resources, and reduced chemical fertilisers and pesticides (1). However, the risk assessment for GM crop methods must be thorough, including molecular characterisation, and food and environmental safety. Only if deemed safe should GM plants be approved for commercial planting and release into the market. Among risk assessments, comprehensive molecular characterisation at the chromosome level is needed, which includes integrated DNA sequencing and determination of the location, number of DNA inserts and flanking sequences, and backbone sequence residues. Furthermore, molecular characterisation information is helpful for selecting GM events yielding high and stable expressed target traits in resulting transgenic offspring lines. (2).

Currently used methodologies to produce GM crops, such as particle bombardment and *Agrobacterium tumefaciens*-mediated transformation, often introduce exogenous DNA into the recipient genome randomly with unintended insertion/deletion or rearrangements, which makes molecular characterisation of GM events highly challenging. In addition, new GM crop events require the development of new, accurate, cost-effective techniques and methods for molecular characterisation. For molecular characterisation of single GM events, several techniques have been reported including Southern blotting, PCR-based methods including real-time PCR and digital PCR, Sanger sequencing, and next-generation sequencing (NGS). (3) In particular, Southern blotting and PCR-based methods coupled with Sanger sequencing are legally required and commonly applied approaches for molecular characterisation in GMO risk assessment guidelines of many countries around the world. For example, Southern blotting is used to determine the copy number of exogenous genes and to confirm the presence/absence of vector backbone sequences; PCR-based methods coupled with Sanger sequencing are used to identify exogenous DNA integrated sites and flanking sequences, examples of which include thermal asymmetric interlaced PCR (TAIL-PCR), ligation-mediated PCR (LM-PCR), and inverse PCR (iPCR). To date, molecular characterisation of most commercialised GM crop events has been revealed through combinations of these techniques (4, 5). However, these approaches are time-consuming and the obtained results are often incomplete. (6) For example, molecular characterisation of GM soybean GTS 40-3-2 has been amended three times since the event was approved for commercialisation in 1994, and rearrangement of the 3’ *NOS* terminator junction and an unintended 70 bp DNA fragment insertion were revealed (7). One transgenic DNA insertion in GM rice event T1c-19 was missed using Southern blotting and TAIL-PCR. (8)

With the rapid development of high-throughput NGS technology in last decade, whole-genome sequencing (WGS) of a single species can be achieved with high accuracy and at reasonable cost, potentially making comprehensive molecular characterisation more accessible (9). Recent progress has been reported for identifying exogenous DNA insert loci and flanking sequences in GM crops and animals using various NGS strategies, such as paired-end WGS, mate-pair WGS, and target enrichment re-sequencing. To date, paired-end WGS (PE-WGS) has been primarily used for molecular characterisation of GMOs. Zhang et al. identified exogenous DNA insertion loci and flanking sequences in transgenic cows using PE-WGS (10). Wang et al. revealed transgene insertion and flanking sequences in GM rice events (SNU-Bt9–5, SNU-Bt9–30, SNU-Bt9–109, G2-6 and PJKS131-2) (11). Guo et al. reported transgene integration in GM soybean events (MON17903, MON87704, L120, L122, L123, GE-J16 and ZH10) (12), Liang et al., revealed the transgene insertion of GM maize MIR162 (6). Yang et al. established a TranSeq bioinformatics tool for full molecular characterisation of GM rice events TT51-1, T1c-19 and G6H1 (8). The mate-pair WGS (MP-WGS) strategy has also been used and benefits from high efficiency and low requirement for sequencing coverage, except for complicated sequencing library construction. Zhang et al. and Srivastava et al. revealed transgene insertion and flanking sequences in transgenic mice and rice lines with sequencing coverage as low as 5× (13, 14). Two target enrichment re-sequencing strategies using one pre-designed tilling probe panel as baits to specially capture transgene-related DNA fragments were reported for identifying transgene integration in transgenic mice and maize lines, respectively. One strategy using microarray hybridisation, and another using liquid phase hybridisation have also been reported (15, 16). NGS has advantages of high throughput, scalability, and time effectiveness, making it suitable for determining transgenic insertion loci and flanking sequences of GMOs. However, due to the inherent short read length of NGS, deciphering the complete sequence and structural arrangement of entire exogenous DNA integration is challenging, especially for GM plant or animal lines containing large DNA insertions with complex structure.

In order to overcome the bottleneck of complex DNA fragment assembly, third-generation sequencing (TGS) techniques aimed at single-molecule sequencing (SMS) were developed based on PacBio and Nanopore sequencing platforms, which could increase the read length to hundreds of kb at the single-molecule level (17). SMS is considered to be a very effective method for deciphering complex genome structures and filling in the gaps of NGS in reference genome assembly (18). Since 2019, various researchers have demonstrated the possibility of TGS at the whole-genome level for molecular characterisation of transgenic animals and plants (19). Osamu et al. identified a single transgene insertion and confirmed a large genomic deletion during transgene insertion in a transgenic mouse strain (20). Paula et al. reported the molecular characterisation of three GM plant species using MinION at the WGS level (21). Anne-Laure et al. identified genomic insertion and flanking sequences in transgenic drought-tolerant maize line ‘SbSNAC1-382’ using single-molecule real-time (SMRT) sequencing coupled with plasmid rescue (17). TGS techniques proved advantageous for unveiling the full sequence and the entire structure of transgene integration and flanking sequences compared with NGS approaches. However, sequencing volume and cost remain very high, and effective bioinformatics tools are needed for data analysis of long reads.

Regardless of the technology applied for molecular characterisation of GM events, the key points are to accurately capture and sequence DNA fragments containing transgenes at low cost. In the present study, we developed a novel universal <u>L</u>arge Integrated DNA <u>F</u>ragments <u>E</u>nrichment strategy coupled with PacBio <u>Seq</u>uencing (Life-Seq) strategy, and a supporting bioinformatics pipeline. Its performance and applicability for molecular characterisation of GM events were evaluated using various commercialised GM crop events.

## Materials and Methods

### Plant materials and DNA extraction

Seeds from NK603 maize, Mon810 maize, TT51-1 rice, GTS 40-3-2 soybean, and RF2 and RT73 canola GM events were kindly supplied by the developers, and authenticity of these GM events was confirmed by event-specific qRT-PCR in our lab. Non-GM maize, rice, soybean and canola seeds were purchased from a local market and confirmed GM-free in our lab. All GM seeds were paired with corresponding non-GM seeds of specific GM content and ground into powder for LIFE-Seq experiments. A total of seven seed powder samples coded S1 to S7 were prepared, including two GM maize samples (S1, NK603; S2, MON810), two GM rice TT51-1 samples (S3 and S7), GM soybean GTS 40-3-2 (S4), GM rapeseed RT73 and RF2 (S5), and non-GM rice, maize, rapeseed and soybean (S6). Details of the components in each sample are listed in Supplemental **Table S1**. For plant genome DNA extraction, a modified hexadecyl trimethyl ammonium bromide (CTAB) procedure was employed (22), and the quality and quantity of extracted DNA were evaluated using a NanoDrop 1000 UV/vis Spectrophotometer (NanoDrop Technologies Inc., Wilmington, DE, USA) and 1% agarose gel electrophoresis with ethidium bromide staining.

**Table 1.** Statistical analysis of reads from PacBio sequencing.

| Samples | Polymerase reads |  | CCS reads |  |  |  |  |
| --- | --- | --- | --- | --- | --- | --- | --- |
|  | Number | Length (bp) | Number | Length (bp) | Min length (bp) | Max length (bp) | Mean length (bp) |
| S1 | 343,991 | 1,947,749,569 | 17,767 | 126,858,938 | 1,069 | 24,771 | 6,175 |
| S2 | 335,491 | 1,870,145,963 | 17,137 | 121,458,324 | 1,385 | 25,373 | 6,083 |
| S3 | 460,133 | 2,674,723,582 | 23,572 | 176,472,547 | 1,072 | 29,827 | 6,351 |
| S4 | 440,023 | 2592557362 | 22,402 | 171,559,156 | 1,104 | 32,878 | 6,445 |
| S5 | 330,565 | 1926612360 | 16,989 | 128,023,275 | 1,249 | 33,575 | 6,393 |
| S6 | 361,475 | 2032302880 | 17,935 | 130,742,309 | 1,109 | 33,137 | 6,142 |
| S7 | 487,813 | 2841288506 | 25,358 | 188,139,490 | 1,257 | 25,360 | 6,333 |

### Universal tiling probe panel design

We designed a universal panel of tiling oligonucleotide probes for hybridisation-based enrichment of transgenic DNAs present in current and commercialised GM crop events. The universal panel of tiling probes was designed to span GM crop events as much as possible by targeting a constructed transgenic DNA library developed in our lab (23), including promoters, terminators, marker genes, exogenous genes, and plant endogenous reference genes (Supplemental **Table S2**). The tiling probe panel consisted of numerous probes by stacking tiles over the target area, and the probe density was increased in areas difficult to cover. The universal probe panel consisted of 150 overlapping regions and the total size of enrichment sequences in the database was ∼172 kb. All target sequences were covered by the universal probe panel and the overlay results file indicated that the synthesised probe coverage was 98.08% for target regions, with 99.64% of target regions estimated to be within range of the synthesised probes (Supplemental **Table S3**). Tilling probe panel design and synthesis were performed by Roche NimbleGen (NimbleGen/Roche, Madison, USA).

**Table 2.**
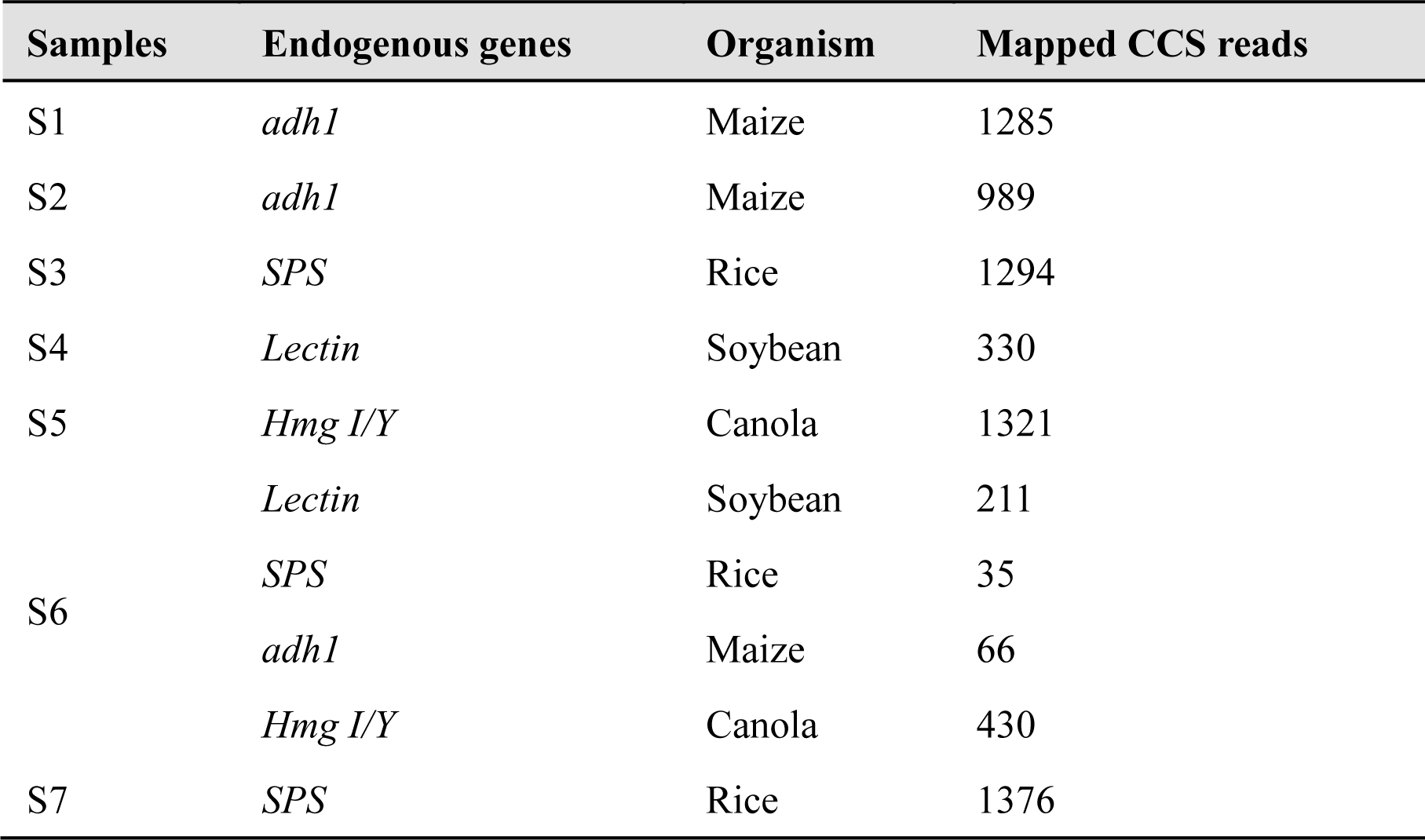
Statistical results for reads mapped to endogenous elements.

**Table 3.**
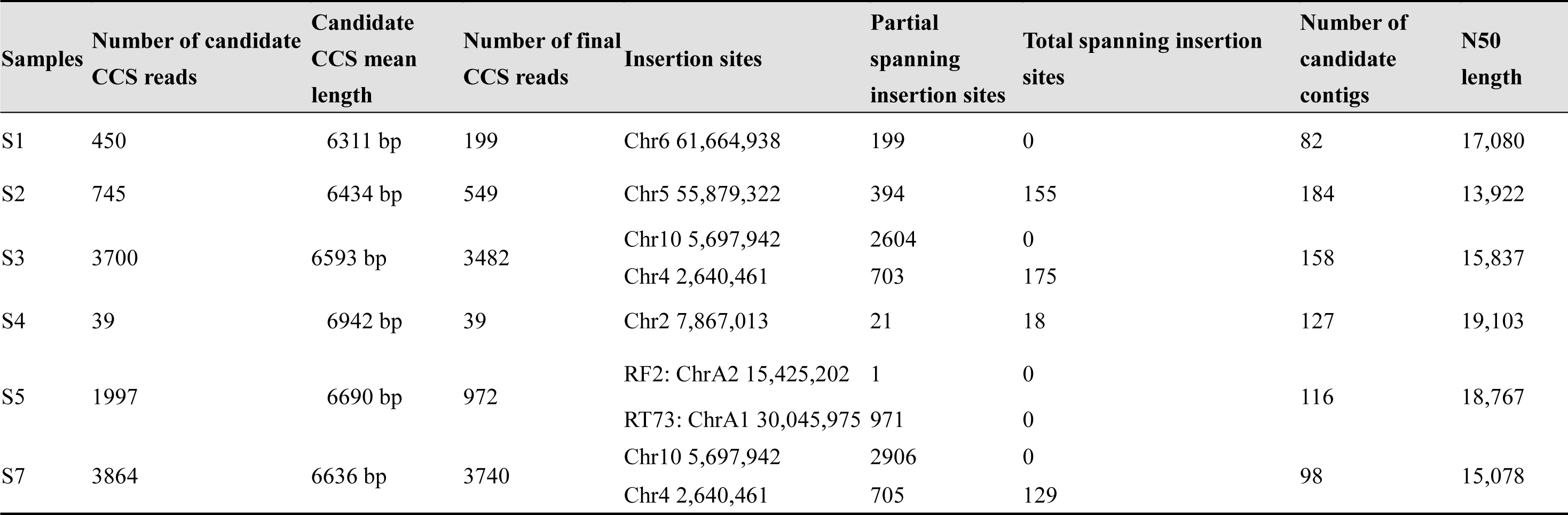
Statistics for candidate CCS reads and contigs covering exogenous DNA integration sites.

### Hybridisation library preparation

For LIFE-Seq, 4 µg samples of genomic DNA were sheared to a peak size of ∼10 kb using a Covaris g-TUBE instrument (Woburn, MA, USA) at 8000 rpm for 60 s. The sheared DNA fragments were end-repaired and deoxyadenosine-tailed in a 60 µL volume with 3 µL End Repair & A-Tailing Enzyme Mix and 7 µL End Repair & A-Tailing Buffer using a KAPA Hyper Prep Kit (KAPA Biosystems, USA) according to the manufacturer instructions (20°C 30 min, 65°C 30 min). For mixed samples, 5 µL of different barcoded adapters were mixed with 10 µL DNA Ligase and 60 µL End Repair and A-Tailing reaction product to a 110 µL total volume and incubated at 20°C for 15 min. A 0.5× ratio of AMPure PB Beads (Beckman Coulter, Inc., USA) was used to clean up the fragments. The DNA fragments were then mixed equally to 2 µg. To narrow the size range of DNA fragments, BluePippin Prep (Sage Science, Beverly, MA, USA) was used with a DNA fragment cut-off of 4.5 kb.

### Large target DNA fragment enrichment

To capture inserted exogenous DNA in GM samples, 6 µL universal probe panel was added to the size-selected DNA fragments and hybridisation buffer and incubated in at 47°C for 16 h. Target DNA capture was performed as previously described (24) with slight modifications using a SeqCap EZ Hybridisation Wash Kit (Nimblegen/Roche, Madison, USA) and Dynabeads M-270 Streptavidin (Thermo Fisher Scientific, USA). Finally, captured DNA fragments were PCR-amplified using Takara LA Taq DNA Polymerase over 15 cycles (Clontech/Takara Bio, Otsu, Japan) then purified using AMPure PB Beads (Beckman Coulter, Shanghai, China). The resulting capture library was quantitated using a Qubit instrument (Thermo Fisher Scientific, USA) and a 2100 Bioanalyzer (Agilent Technologies, USA) to confirm the concentration and size distribution, respectively.

### PacBio SMRTbell library construction and sequencing

A sequencing library for the PacBio platform was generated after ligating PacBio universal primer (GCAGTCGAACATGTAGCTGACTCAGGTCAC, 100 µM, TE pH 8.0; Integrated DNA Technologies, USA). A Template Prep Kit (PacBio, USA) was used to remove failed ligation products. Purification was achieved using a 0.8× ratio of AMPure PB Beads (PacBio, USA). The final sequencing library was validated using a 2100 Bioanalyzer High-sensitivity DNA chip (Agilent Technologies, USA) and a Qubit 2.0 Fluor-meter High-sensitivity Kit (Life Technologies, USA). The final sequencing library was sequenced using a Sequel Sequencing Kit 3.0 and an SMRT Cell following the manufacturer’s protocol.

### Analysis of LIFE-Seq data

PacBio sequencing raw datasets were demultiplexed based on different barcodes and adapters were removed using lima (https://github.com/pacificbiosciences/barcoding). The PacBio SMRT Portal workflow (v3.4.1) was used to trim Circular Consensus Sequencing (CCS) reads (https://ccs.how/how-does-ccs-work.html). The trimmed CCS reads were mapped to reference genomes (rice genome GCF_000005425.2; soybean genome GCF_000004515.5; maize genome GCF_000005005.2; rapeseed genome GCF_000686985.2) or transgenic DNA sequences using Minimap2 (25) and visualised by Integrative Genomics Viewer (IGV) (26). Both mapped CCS reads were imported for transgene integration site(s) analysis. CCS reads were assembled by Flye 2.8.2 (https://anaconda.org/bioconda/flye). The assembled contigs were then imported for exogenous DNA insertion structure analysis using NCBI BLASTN (https://blast.ncbi.nlm.nih.gov/Blast.cgi).

## Results

### The main LIFE-Seq workflow

To comprehensively decipher exogenous DNA integration (integration sites, entire inserted DNA sequences and structures, and flanking sequences), LIFE-Seq was developed. LIFE-Seq consists of four major procedures: universal tilling probe design, long target DNA enrichment, PacBio sequencing, and data analysis (**Figure 1**).

**Figure 1.**
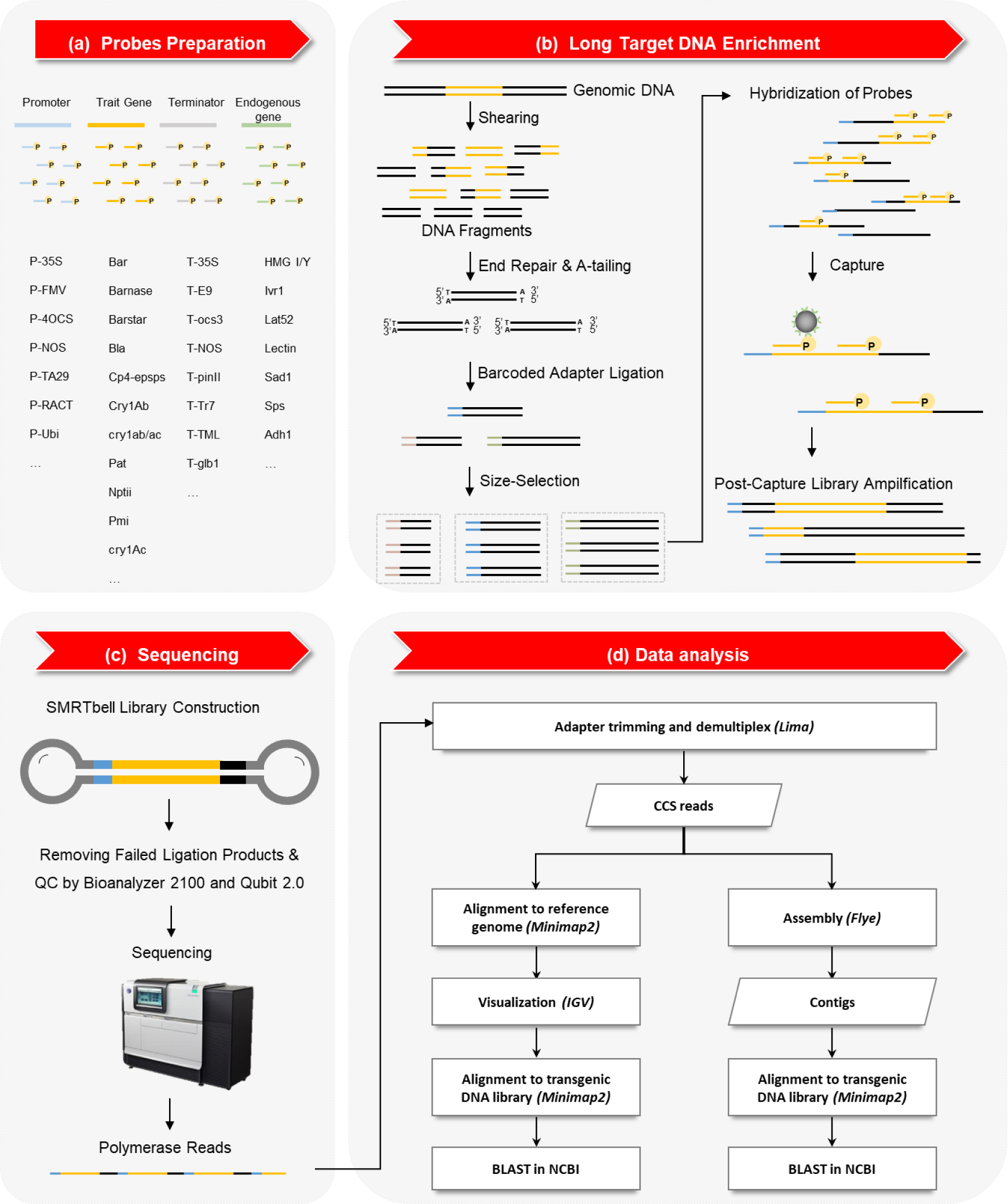
Schematic diagram of the LIFE-Seq approach. The method includes four main steps: universal tiling probe panel design, long target DNA fragment enrichment, PacBio sequencing, and sequencing data analysis.

i. In universal tilling probe design, we aimed to design a universal tilling probe panel targeting as many GM events as possible, rather than one probe panel targeting a single GM event. Therefore, we constructed a transgenic DNA library for tilling probe design based on our previously developed database, including commonly used transgenic elements (promoters, terminators, marker genes and exogenous genes) and endogenous reference genes of major crop species (**Figure 1a**).
ii. Long target DNA enrichment consisted of several experimental operations, including genome DNA random fragmentation, end repair, A-tailing ligation, barcode adapter ligation, size selection, tiling probe hybridisation and capture, and long-range PCR (**Figure 1b**).
iii. In PacBio sequencing, the enriched target DNAs were ligated with PacBio blunt to construct the SMRTbell sequencing library, and loaded onto a cell using the PacBio Sequel system for long read-length sequencing (**Figure 1c**).
iv. In data analysis, a bioinformatics pipeline for exogenous DNA integration analysis was established (**Figure 1d**). Raw polymerase reads were trimmed and demultiplexed by lima. CCS reads were firstly aligned against the corresponding plant reference genome, and matched CCS reads were visualised using IGV. Next, matched CCS reads were aligned against transferred plasmid sequences, and matched CCS reads were filtered and BLAST searched against NCBI databases to analyse integration structures. Meanwhile, contigs were generated and aligned against the transgenic DNA library. Matched contigs were used to confirm exogenous DNA integration loci and integrated structures following BLAST searches.

### PacBio sequencing data generation from tested samples

A total of seven samples (S1 to S7) from four crop species were designed and tested. The length of enriched DNAs in each sample was between 5 kb and 10 kb with a peak of ∼6298 bp (Supplemental **Figure S1**). PacBio sequencing data or enriched DNAs from all seven samples are shown in **Table 1**. A total of 1.86 Gb, 1.78 Gb, 2.55 Gb, 2.47 Gb, 1.84 Gb, 2.7GG, 1.94 Gb and 2.71 Gb, corresponding to 343,991, 335,491, 460,133, 440,023, 330,565, 361,475 and 487,813 polymerase reads were acquired for S1 to S7, respectively. After filtration, 17,767, 17,137, 23,572, 22,402, 16,989, 17,935 and 25,358 CCS reads, corresponding to S1−S7, were retained for further analysis. The length of CCS reads ranged from 1069 bp to 33,575 bp, and the average length of CCS reads ranged from 6083 bp to 6445 bp in the seven samples. Statistical analysis of the sequencing data showed that the distribution of enriched DNAs were similar in length and quantity, regardless of the crop species tested (Supplemental **Figure S2**).

### Evaluation of LIFE-Seq performance using plant endogenous reference genes as calibrators

In the designed universal tiling probes panel, the sequences of endogenous reference genes of various crop species were used as calibrators to evaluate the performance of LIFE-Seq, and used as positive controls in practical sample analysis. In all seven samples tested, thousands of CCS reads containing entire or partial plant endogenous genes were successfully enriched and sequenced (**Table 2**), and the assembled contigs with an average length of 23,756 bp covering the corresponding endogenous reference genes were obtained from the CCS reads. For example, 1294 CCS reads mapped to the rice endogenous reference gene *SPS* in S3, generating a contig of 23,894 bp. Sample S7 was designed as a repeat of S3, 1376 CCS reads mapped to the rice *SPS* gene, and a contig of 31,222 bp covering the *SPS* gene was assembled, which was exactly the same as that for S3. In the negative control sample of S6 comprising a mixture of non-GM soybean, maize, rice and canola, 742 CCS reads were obtained, including 211 CCS reads for the soybean *Lectin* gene, 66 CCS reads for the maize *adh1* gene, 35 CCS reads for the rice *SPS* gene, and 430 CCS reads for the canola *HMG I/Y* gene, generating 166 assembled contigs with an N50 length of 17,190 bp. All corresponding endogenous reference genes were observed in the assembled contigs. In the other four samples, the obtained CCS reads and contigs successfully covered the corresponding endogenous reference genes. These results showed that all target DNA fragments of plant endogenous reference genes were successfully enriched and sequenced through LIFE-Seq, indicating that no obvious bias in target enrichment, even though the universal tiling probe panel consisted of thousands of probes. We believe that LIFE-Seq employing the universal tiling probe panel is effective and unbiased for large target DNA fragment enrichment from tested samples.

### Characterising exogenous DNA integration of various GM crop events using LIFE-Seq

According to the pipeline shown in **Figure 1**, the sequenced data from the seven samples were carefully analysed. CCS reads mapped to both sequences of plant reference genomes and the transgenic DNA library were selected as candidates to further characterise exogenous DNA integration, including the complete sequences and structures of the whole insertions. The number of CCS reads covering the exogenous DNA integration sites for each sample are listed in **Table 3**. The elements from the transgene DNA library included in the candidate CCS reads were aligned and statistically analysed, and the distributions of the elements are shown in **Supplemental Figure S3**. The results indicate that both the expected exogenous DNAs and plant endogenous reference genes were enriched and sequenced with high efficiency and accuracy, although the number of CCS reads mapped to plant endogenous genes was higher than the number mapped to transgenic elements.

#### S1 (NK603 maize)

The NK603 event reported by Monsanto Company was achieved through particle bombardment transformation, resulting in the expression of 5-enolpyruvylshikimate-3-phosphate synthase (EPSPS), allowing the plant to survive the otherwise lethal application of glyphosate (27). The S1 sample comprised seeds of GM maize event NK603 with a GM content of 50.0%. In the S1 tests, 17,767 CCS reads were obtained from polymerase reads for further analysis of exogenous DNA integration. After mapping the 17,767 CCS reads to the transgenic DNA library and maize reference genome, 450 candidate CCS reads were screened out.

Among the 450 CCS reads, 199 reads (55.78%) were considered positive, covering the partial T-DNA integration site after careful checking of the mapping results for the transgenic DNA library, including five CCS reads spanning upsteam of the integration site and 194 reads spanning downstream (**Table 3**). However, none of the individual CCS reads covered the entire exogenous DNA insertion, which might be because the length of enriched fragments was slightly shorter than that of DNA insertions. IGV visualisation and BLASTN analysis revealed the exogenous DNA located at the Chr6 61665252 locus (**Figure 2a**). A total of 82 contigs were assembled from 17,767 CCS reads, including one contig covering the entire exogenous DNA insertion and its flanking sequences. BLASTN analysis showed that 7,462 bp of exogenous DNA was inserted into the recipient maize genome at Chr06 61665252 with a 3 bp deletion, and the adjacent 1752 bp upstream and 400 bp downstream flanking sequences. The entire structure of the insertion is shown in **Figure 3**, including the rice *Actin* promoter, rice Actin 1 intron, CTP2, Cp4-epsps, NOS terminator, enhanced CaMV35s promoter, hsp70, CTP2, Cp4-epsps, NOS terminator, partial rice actin promoter, and partial rps11 and rpoA genes, in this order.

**Figure 2.**
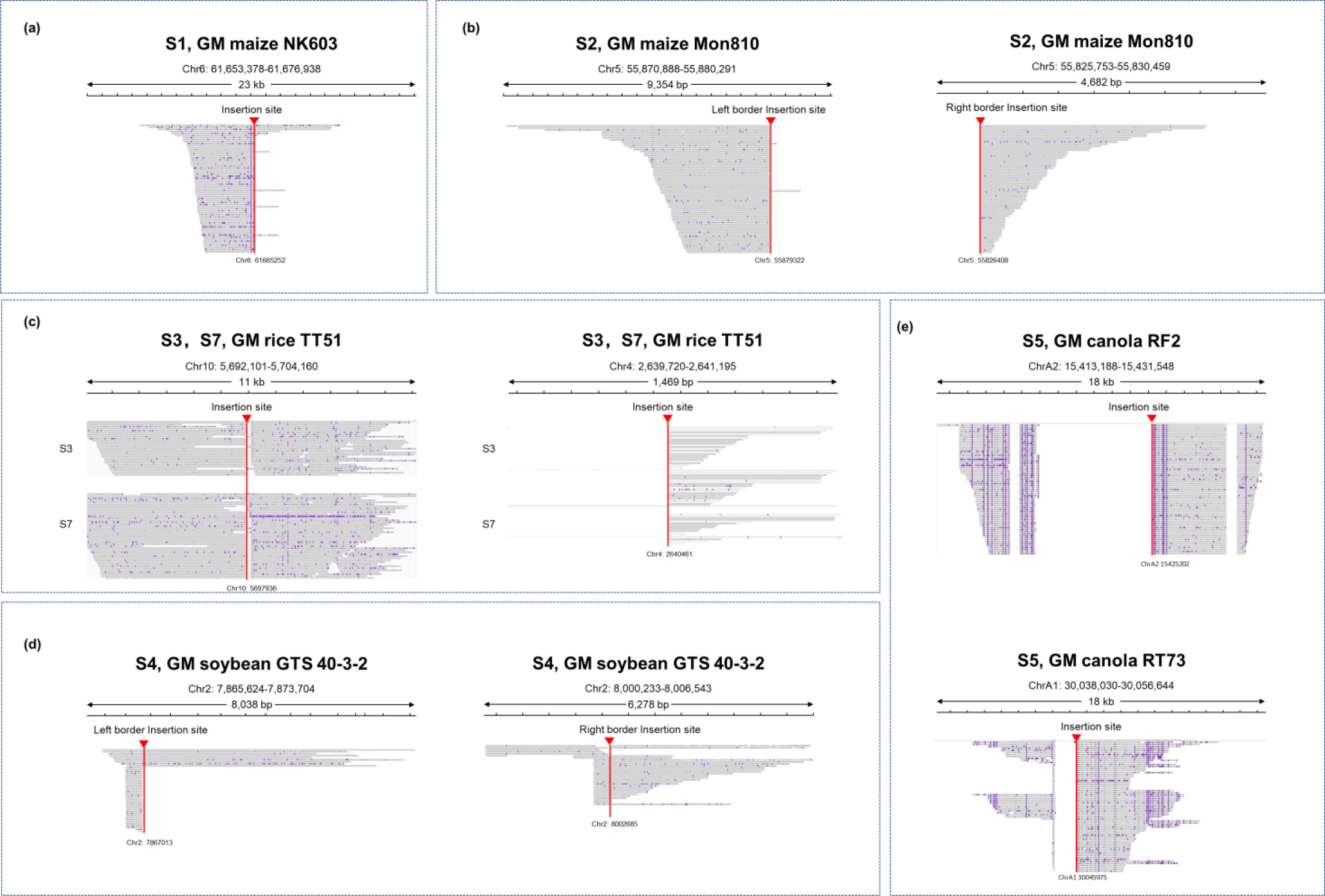
IGV alignment of obtained CCS reads for tested samples in 1–23 kb windows spanning the T-DNA insertion sites. The insertion sites are visible as sharp vertical read lines.

**Figure 3.**
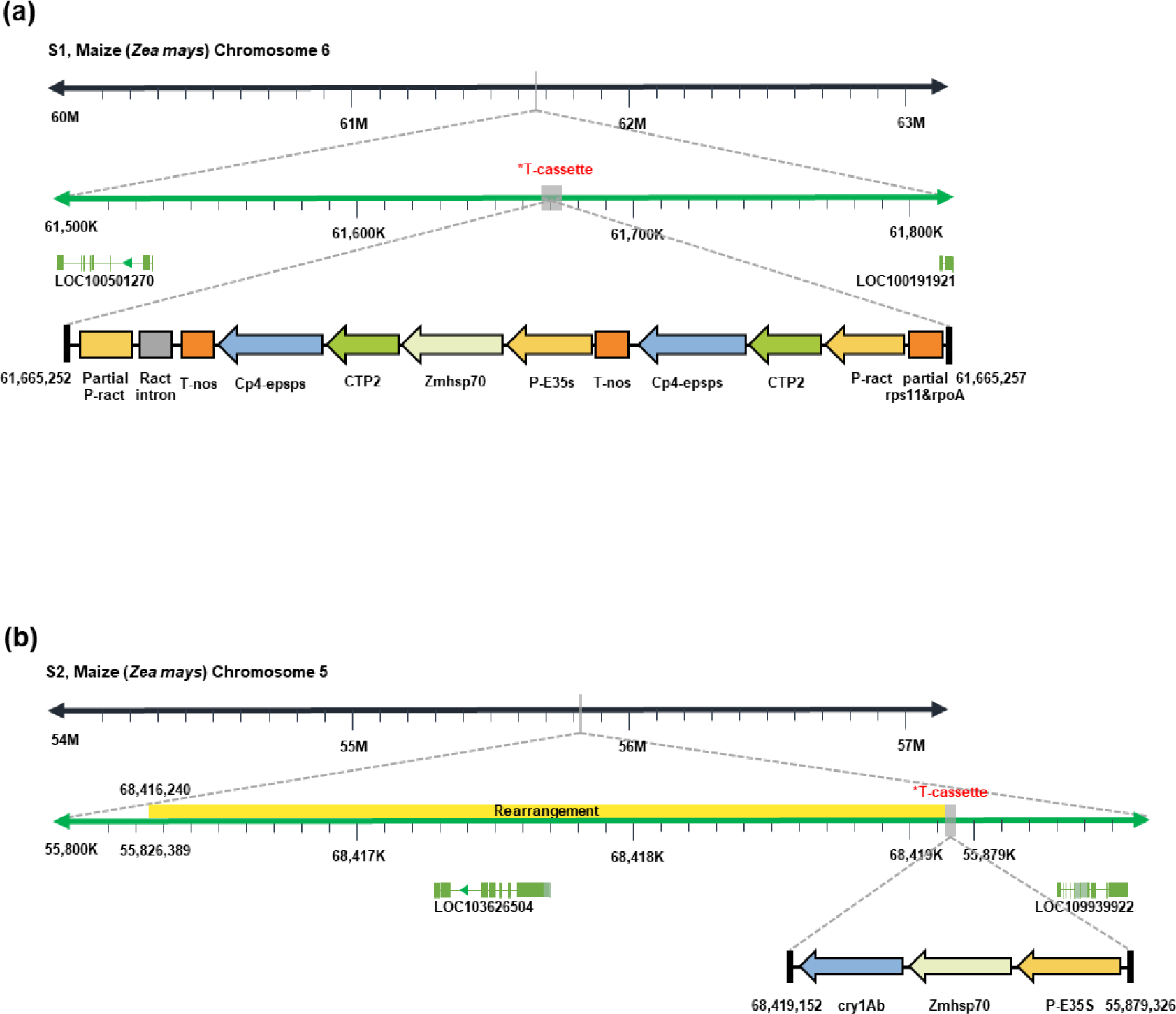
Schematic diagram of the whole structure and arrangement of transgene integration in GM maize event NK603 (sample S1) and MON810 (sample S2).

Bioinformatics analyses of the junctions between the original genome and inserted sequences did not reveal any potential creation of unintended genes or any disruption of pre-existing genes in the recipient maize genome. The obtained structures and flanking sequences of exogenous DNA integration were consistent with previously reported data from Southern blotting, genome walking, and Sanger sequencing analyses (28). The assembled contigs resulting from LIFE-Seq further confirmed the previous results based on complete sequence information. In addition, the assembled results also included another two contigs with lengths of 23,584 bp and 17,068 bp, which were exactly the same as the regions covering maize *adh1* and *hsp70* genes in maize genome, respectively. This indicates that LIFE-Seq was highly credible and accurate because tiling probes for *adh1* and *hsp70* were also included in the universal tiling probe sets.

#### S2 (MON810 maize)

The S2 sample comprised GM maize MON810, produced by inserting the *cry1Ab* gene cassette related to insect resistance. Data analysis showed that 745 CCS reads matched both the transgenic library and the maize genome. Among the 745 CCS reads, 549 CCS reads (73.69%) were classified as candidate reads covering the partial or whole exogenous DNA integration sites, including 394 CCS reads spanning partial exogenous DNA integration sites and their adjacent sequences, and 155 CCS reads covering the whole DNA integration sites and their adjacent sequences (Table 3). IGV visualisation indicated exogenous DNA integration at maize locus Chr05 (**Figure 2b**).

After assembly, 184 contigs were generated from all CCS reads. The BLASTN results showed that contig 163 with a length of 14,016 bp consisted of four pieces of DNA from 55,874,135 to 55,879,326 of maize chromosome 5, 3530 bp of exogenous DNA, 68,416,240 to 68,419,152 of maize chromosome 5, and 55,826,389 to 55,827,211 of maize chromosome 5, indicating exogenous DNA integration at locus 55,879,326 of chromosome 5.

The results also revealed transgene insertion in the maize zein gene cluster with the rearrangement of both exogenous DNA elements and maize genome DNA during transgene transformation. The inserted exogenous DNA was made up of the *CaMV35s* promoter, *Zmhsp70* intron, and truncated *Cry1ab* gene in this order, which is obviously different from the transferred plasmid PV-ZMBK07. Only the partial expression cassette of the *Cry1ab* gene was integrated into the maize genome, and the 3’ end of *Cry1ab* and the *NOS* terminator were lost during integration. The maize Zm-upl gene from 68,416,240 to 68,419,152 of maize chromosome 5 was rearranged and inserted after exogenous DNA integration into the locus at 55,879,302 of Chromosome 5. In previous reports, including those from the developer and the Agricultural Research Center in Belgium, a 3’ truncated *Cry1ab* cassette was inserted into MON810 maize and its 5’ adjacent junction sequence was confirmed, and the exact 3’ limit of the *Cry1ab* gene and its adjacent host DNA is unknown (29). Previously reported molecular data do not completely fulfil the EU and China requirements for molecular characterisation, although the MON810 event has been commercialised in plants for more than 20 years (30). Our S2 sample analysis is the first full molecular characterisation of the MON810 event using LIFE-Seq, revealing the complete structures and sequences of transgene integration, confirming previous speculation about the rearrangement of transgenic integration in the MON810 event.

#### S3 and S7 (TT51-1 rice)

GM rice TT51-1 possessing the insect resistance trait was produced by microparticle bombardment with two vectors (pFHBT1 and pGL2RC7) simultaneously (31). Two parallel samples (S3 and S7) containing 50.0% GM rice TT51-1 were prepared and tested to explore exogenous DNA integration and evaluate the reproducibility of LIFE-Seq.

For the S3 sample, 3,700 candidate CCS reads were obtained, which were mainly divided into two groups according to rice chromosome loci; one group of candidates comprising 2,604 reads clustered on chromosome 10 and the other (878 reads) clustered on chromosome 4 (**Table 3**). All candidate reads were visualised by IGV, which indicated two exogenous DNA integration sites in rice event TT51-1; one located around Chr10: 5,692,101, and the other located around Chr04: 2,639,720 (**Figure 2c**). In the group with 2,604 CCS reads, all reads consisted of partial exogenous DNA and partial rice genome fragments, and none covered the entire exogenous DNA insertion. In the other group (Chr04) with 878 CCS reads, 175 reads covered the whole exogenous DNA insertion, and the other 703 reads covered partial exogenous DNA insertion. The 175 CCS reads included a truncated *Cry1Ab/Ac* gene cassette of 383 bp inserted into rice Chr04 at 2,639,720, including *P-ract* and *Cry1Ab/Ac* (**Figure 4**). The *de novo* results for all CCS reads yielded 158 contigs with an N50 of 15,837 bp in length. Sequence analysis showed that Contig 15 with the length of 15,910 bp consisted of 8993 bp of rice genome DNA and 6917 bp of partial integrated exogenous DNA, indicating an exogenous DNA integration located at locus Chr10 5,697,935 (**Figure 4**). After manually assembling CCS reads, the whole insertion was determined. Contig 84 with a length of 17,710 bp was the same as the rice *Actin* gene region located at Chr03 29,074,005 of the rice reference genome, consistent with the use of the rice endogenous *Actin* promoter in the exogenous gene expression cassette to generate TT51-1. Compared with previously reported results from PE-WGS and genome walking analyses (8, 32), we confirmed two rearranged exogenous DNA integrations in Chr10 and Chr04 with complete sequences in GM rice TT51-1, including a 13 bp deletion at the Chr10 insertion site and a filled rice DNA fragment in the truncated *Cry1Ab/Ac* cassette at the Chr4 insertion. However, several short independent contigs matching partial exogenous DNA integration were obtained in PE-WGS analysis (8).

**Figure 4.**
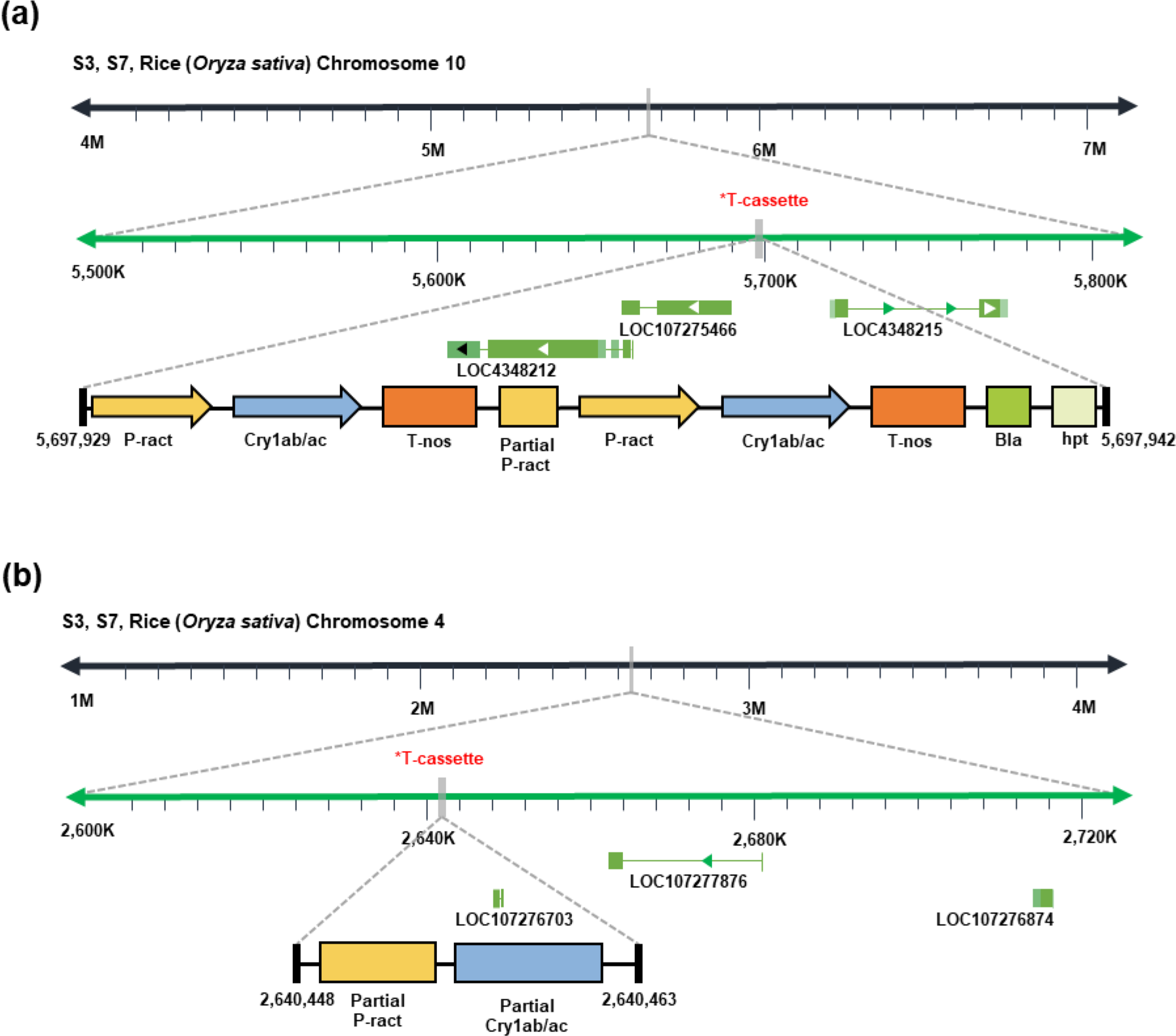
Schematic diagram of the whole structure and arrangement of transgene integration in GM rice event TT51-1 (samples S3 and S7).

For the S7 sample, 487,813 CCS reads were obtained, and 3,864 candidate CCS reads were identified as candidate reads. Among the 3,864 candidate reads, 2,906 CCS reads clustered around Chr10: 5,692,101, and the other 705 CCS reads clustered around Chr4: 2,639,720 (**Table 3**). *De novo* analysis resulted in 98 contigs, including contigs containing exogenous DNA insertions and the rice *Actin* gene, which was highly similar to the results for the S3 sample. Both the results from the candidate reads and *de novo* analysis of S3 and S7 samples yielded the same molecular characterisation data for TT51-1 rice, demonstrating that the developed LIFE-Seq method has very high reproducibility.

#### S4 (GTS 40-3-2 soybean)

GTS 40-3-2 GM event soybean plants, related to herbicide tolerance, were transformed with the cloning vector PV-GMGT04 by microparticle bombardment (7). Sample S4 containing 5.0% GM soybean GTS 40-3-2 was prepared and used to evaluate the applicability of Life-Seq to low GM content samples. This yielded 39 candidate CCS reads from total 440,023 reads, including 18 CCS reads involving complete exogenous DNA insertion (Table 3). The sequences of the 18 CCS reads revealed exogenous DNA integration at the Chr02 7,867,013 locus with a length of 2420 bp, including elements of the CaMV35S promoter, the cp4-epsps gene, the NOS terminator, and partial cp4-epsps (248 bp) in a tandem arrangement (**Figure 2d**, **Figure 5**). A total of 127 contigs were obtained from *de novo* analysis, and the BLASTN search results identified Contig 62 with a length of 8621 bp following exogenous DNA integration, including the complete exogenous DNA insertion of 2420 bp. The sequence of Contig 62 also contained soybean genome DNA fragments derived from the Chr2 68,416,243 68,419,146 loci, hence the rearrangement likely occurred in the soybean recipient genome during transgene insertion, as reported previously by Monsanto (**Figure 5**) (7).

**Figure 5.**
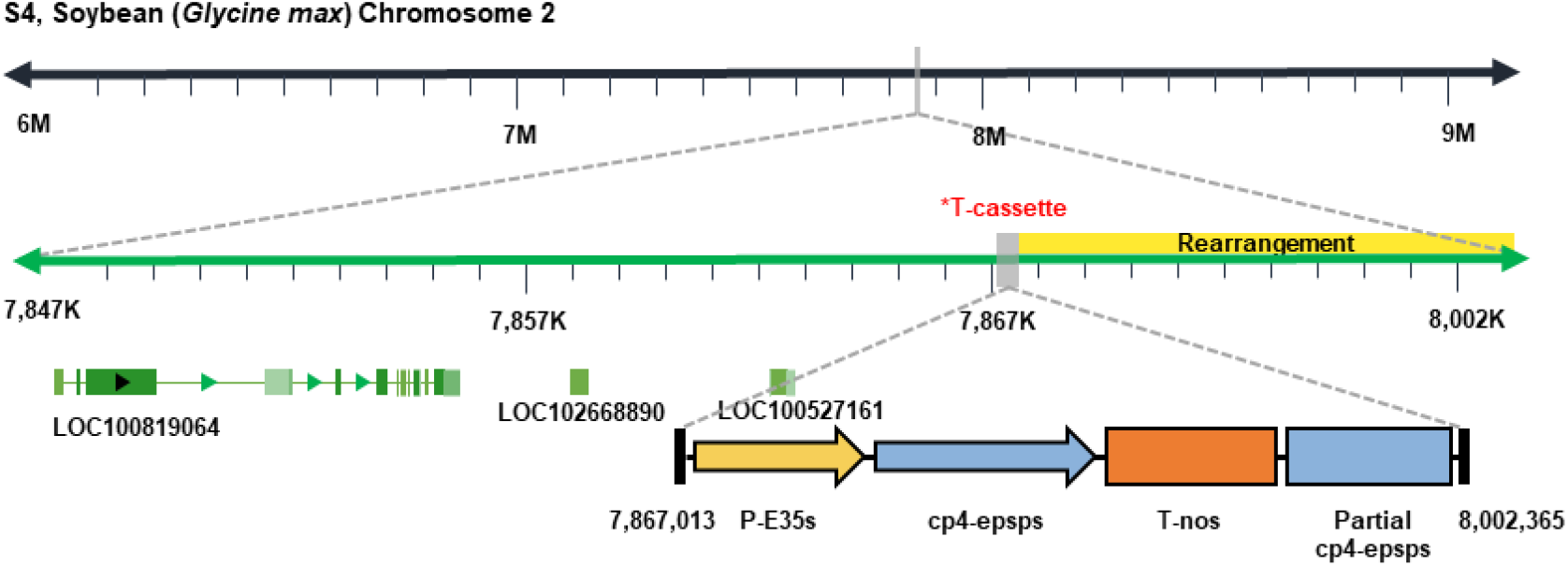
Schematic diagram of the whole structure and arrangement of transgene integration in GM soybean event GTS40-3-2 (sample S4).

Contig 48 with a length of 20,965 bp was the same as the *Lectin* gene region of the soybean reference genome, which also indicated that LIFE-Seq performed with high credibility and accuracy. The results for S4 also showed that LIFE-Seq has high sensitivity, and could be used to characterise exogenous DNA integration in samples with low GM content.

#### S5 (RT73 and RF2 rapeseed)

Sample S5 was a mixture of with GM canola RT73 and GM canola RF2 with the mass ratio of 99:1, and this was used to evaluate the applicability of LIFE-Seq for mixed samples with GM content as low as 1.0%. Canola RT73 included two introduced genes (*CP4 EPSPS* and *gox247*) that confer herbicide tolerance. Canola RF2 was developed via *A. tumefaciens*-mediated infection with pTVE74RE to restore fertility. After data analysis, 1997 CCS reads were obtained and used to further characterise exogenous DNA integration (Table 3). IGV visualisation showed that all candidates were clustered into three groups; the first with 971 reads comprising exogenous DNA insertion of RT73, the second with only one read spanning exogenous DNA insertion of RF2, and the third with 996 reads covering the rapeseed endogenous *HMG I-Y* gene sequence. Further analysis showed that all 971 reads covered the partial exogenous DNA insertion and its flanking sequences without complete transgene integration, including 29 reads and 942 reads covering the partial 5’ and 3’ ends of transgene integration, respectively (**Figure 2e**). The full sequence of transgene integration was obtained by assembling these reads. The results showed that the transgene was inserted into the rapeseed gene at ChrA01 30,045,975 with a length of 6224 bp DNA in RT73, including the FMV35s promoter, gox247, E9 terminator, FMV35s promoter, Ctp2, Cp4EPSPS, and E9 terminator in this order (**Figure 6a**). The flanking sequences of the integration site included an unpublished rearrangement of genomic DNA ranging from 33,702,066 to 33,703,177 upstream of the insert site in the rapeseed genome. BLASTN analysis of the single read with a length of 5722 bp showed that the transgene was inserted into the rapeseed genome at the ChrA02 15,425,202 locus in RF2, including the 5’ flanking sequence of transgene insertion and the exogenous NPTII gene cassette regulated by the NOS promoter and the ocs 3 terminator (**Figure 6b**). The identified insertion site and flanking sequence were the same as previously reported (33).

**Figure 6.**
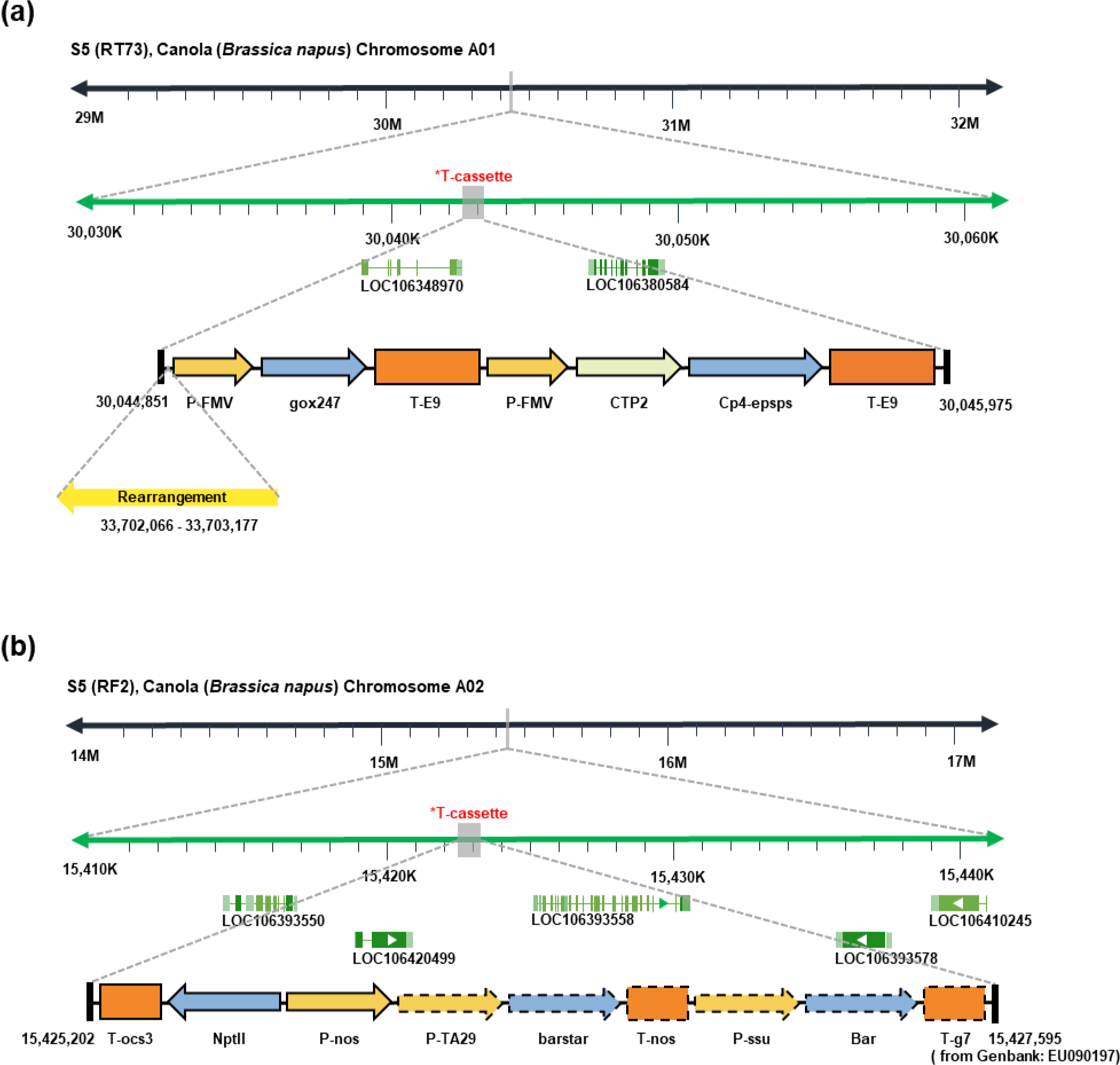
Schematic diagram of the whole structure and arrangement of transgene integration in GM canola RF2 and RT73 (sample S5).

*De novo* analysis of all CCS reads generated 116 contigs. The BLASTN results showed that Contig 43 with a length of 20,634 bp covered the entire integration of RT73, and consisted of 14,349 bp of rapeseed genomic DNA and 6285 bp of inserted transgene DNA. The sequence of Contig 43 was the same as those derived from the above candidate CCS reads. None of the contigs included transgene insertion of RF2, indicating that RF2 information was missed in the direct *de novo* analysis of all CCS reads.

#### S6 (non-GM samples)

Sample S6 containing four non-GM crop species served as a negative control in LIFE-seq experiments, and 792 CCS reads mapped to the transgenic DNA library. Further analysis showed that all CCS reads included native endogenous genes of corresponding species, and no transgene-related exogenous DNA integration was observed (Supplemental **Figure S4**). For example, 431 CCS reads mapped to the maize HMG gene, 212 CCS reads mapped to the soybean Lectin gene, and 16 CCS reads mapped to the rice actin promoter.

## Discussion

Molecular characterisation is crucial for GM event analysis for risk assessment and approval for commercialisation. Although GM crops have been approved and planted for more than 20 years, molecular characterisation data for approved GM events is continually advancing as NGS and TGS technologies are applied. However, full molecular characterisation of GMOs with high accuracy and low cost remains a major goal. In the present study, we developed a novel strategy named LIFE-Seq which combines long target DNA fragment enrichment and PacBio single molecule sequencing for molecular characterisation of GMOs. Compared with NGS-WGS and TGS-WGS strategies (8, 34), LIFE-seq has advantages including strong universality, low cost, easy data processing, high data integrity and accuracy, and high sensitivity.

For LIFE-Seq, a universal tilling probe set was designed to target 150 commonly used transgenic elements, exogenous genes, and crop endogenous reference genes. This design makes the universal tilling probe set suitable for most commercialised GM events with a theoretical coverage of 99.09%, hence the developed LIFE-seq method should be suitable for molecular characterisation of 325 out of 328 commercialised events. Our results also confirmed that the target DNA fragments were enriched successfully from the primary GM crop species (rice, soybean, maize and canola) and GM events (TT51-1, GTS 40-3-2, MON810, NK603, RT73 and Rf2) using the universal tilling probe set, even for RF2 with GM content as low as 1% in mixed sample S5. The target enrichment strategy can be coupled with NGS for the detection of genomic structural variants, copy number variation, and new gene identification, including whole exome sequencing (WES) and Southern by sequencing (SBS) (35). However, the tilling probe set design is often specific for a single species in WES (36), and/or specific to the transformed plasmid for each GM event in SBS (35). In the present work, we broadened the application scope of the tilling probe set to most GM crop species and GM events based on the complete transgenic DNA library, with one probe set for all GM events with strong universality. For all seven samples tested, only one tiling probe set was used to enrich all target DNA fragments, and one mixed sequencing library was constructed using different barcodes and sequenced in one lane, which reduced the cost dramatically compared with SGS and TGS at the whole-genome level for analysis of GM samples individually.

In current NGS approaches, data analysis is key and depends on well-designed pipelines and professional bioinformatics software due to the short read length (≤ 250 bp). For molecular characterisation of GMOs, although some programs and pipelines have been developed (13, 37), testing practical samples remains complicated since there is a need to fine-tune the procedure when analysing varied GM events. However, the length of obtained CCS reads is often longer than 5 kb in LIFE-Seq, which decreases the requirements for short sequence assembly and reduces the difficulty of data analysis caused by complex structures. Therefore, the data analysis process is much simpler than that of NGS. In LIFE-Seq, two different data process pipelines were developed; one for selecting reads covering the whole or partial exogenous DNA integration, and another for the assembly of all CCS reads to identify contigs covering the whole DNA integration. Both pipelines are much easier to use than that of NGS, although our results showed that the first pipeline performed better and was easier to use than the second pipeline. In this work, CCS reads containing the entire exogenous DNA integration were obtained for MON810, GTS40-3-4 and TT51-1 GM events. From these reads, information on exogenous DNA insertion site, flanking sequence, copy number, and the entire structure were obtained without further data analysis. Even for the NK603 and RT73 samples, the whole exogenous DNA integration could be easily determined by simply splicing only two selected CCS reads.

During the process of GM crop production, transgenes are introduced into the recipient genome randomly, and sometimes accompanied by host gene interruptions, multiple exogenous DNA insertions in one or more chromosomal locations, and host genome rearrangement around the insertion site, which may affect the functional expression and genetic stability of trait-determining genes (38). Therefore, comprehensively characterising the whole transgene integration in the recipient genome is a prerequisite not only for GM crop production, but also for commercialisation. Although PE-WGS has been used successfully for molecular characterisation of GMOs, especially for the identification of integration sites and flanking sequences (13, 39), the short read length is still a bottleneck in NGS, which limits its application for providing the ‘overall picture’ of transgene integration, such as the entire sequence and structure of complex integrations involving host endogenous genes and/or repetitive regions, multiple transgene insertions in one or more chromosomal locations, host genome rearrangement around integration sites, and repetitive regions of GMOs (17). LIFE-Seq provides an effective solution for specifically describing transgene integration, and the results for four mainly GM crop events proved that the comprehensive information on transgene integration was highly accurate and reliable, even for samples with GM content as low as 5%. Among the six GM samples, transgene integration of five GM events (NK603, MON810, TT51-1, GTS40-3-2 and RT73) was fully deciphered using LIFE-Seq, but not for the RF2 event in mixed sample S5 with a very low GM content of 1.0%. Although these GM events have been commercialised and plants grown for more than 20 years, the full sequences of inserted exogenous DNA and adjoining regions are reported in their entirety for the first time in the present work. Further analysis of the revealed sequences and structure of the transgene integrations of these events can provide novel findings. For example, in Mon810, GTS 40-3-2 and RT73 GM events, host genome rearrangement around the integration site was observed, which was not observed or confirmed in previous reports (33, 40).

In currently used approaches, host genome rearrangement is difficult to detect due to the limitation of short read length. For example, short DNA fragments are mainly identified using genome walking coupled with Sanger sequencing or NGS, while isolated flanking regions of insertion sites are often limited to <1 kb in length, hence rearrangements may be overlooked. A liquid chip capture strategy incorporating NGS has been used to explore transgene elements and constructs, but only a few incomplete integrations were observed through single *de novo* analysis (33). Amanda et al., also developed a microarray hybrid capture and NGS approach, and revealed more unexpected transgene insertion junctions and complex sequences of transgenes, but failed to decipher the entire structures and full sequences of integrated transgenes (41). The results from the five GM events the four crops showed that LIFE-Seq can decipher the entire configuration of transgene insertion with higher integrity and accuracy than previously reported targeted capture strategies.

A total of six practical GM samples were tested in LIFE-Seq, including two duplicates (S3 and S7; TT51-1), one low GM content sample (S4; 5% GTS 40-3-2) and one mixed sample (S5; 99% RT73 and 1% RF2). The results of duplicate experiments showed that two exogenous DNA integrations, in Chr04 and Chr10, were observed in S3 and S7, and the full sequences and complete structures of transgene insertions were identical, indicating that LIFE-Seq achieves good reproducibility, although the designed probes and DNA hybridisation were easily influenced by GC content and secondary structure variation among different DNAs. The results also showed that biological replicates are not necessary for LIFE-Seq.

For molecular characterisation of GM events, pure GM material is generally needed, and few studies have explored the possibility of characterising mixed samples with low GM content since the collection of exogenous DNAs with sufficient coverage is challenging (42). In the S4 sample with 5% GM soybean GTS 40-3-2, full molecular characterisation was achieved, including the full sequence of exogenous integration and rearrangement in the recipient genome. In the S5 sample comprising a mixture of 99% RT73 and 1% RF2, full molecular characterisation of RT73 was achieved and a novel rearrangement around the integration site was identified, while partial exogenous DNA integration was observed in RF2. These results indicate that LIFE-Seq is suitable for molecular characterisation of mixed GM samples with GM content as low as 5%. Furthermore, partial sequences and structures of transgene integration could be detected in samples with a GM content of only 1%. In previously reported target capture and NGS approaches, it is almost impossible to achieve molecular characterisation of mixed GM events due to the short read length, which limits the distinguishability of similar transgene elements, tandem repeat sequences, and inversion sequences from multiple GM events. The results from S4 and S5 samples also showed that LIFE-Seq is applicable for mixed samples and/or low GM low content samples, although the full configuration of exogenous DNA insertion was not successfully determined for S5 containing 1% RF2.

Although the developed LIFE-Seq method has many advantages, there is still room for further improvement. The universal tilling probe set could be extended to cover more transgene elements, and potentially exogenous trait-related genes, to expand its detection scope for unknown GM events. Optimisation of large DNA fragmentation and DNA hybridisation might also be useful whole exogenous DNA integration. The target capture strategy based on DNA fragments binding to nucleases or specific proteins could also be employed, which might improve the efficiency of large DNA enrichment, such as dead Cas9, dead Cpf1, and dead pAgo (43). The Oxford Nanopore MinION single-molecule sequencing platform could also be coupled with large DNA enrichment, which could potentially generate long read sequences rapidly (44).

However, obtaining CCS reads requires the sequencing of insertion fragments more than three times, the total length of sequencing is limited, self-connected joints or unidentified barcodes may be problematic, and the number of CCS reads in exogenous fragments is small, which may lead to insufficient data to characterise the assembly process. Additionally, splicing software may not be able to provide full-length splicing results.

## Acknowledgements

The authors would like to thank Mr. Yang Lv and Mr. Jie Zong for their kindly comments on bioinformatics analysis. They also thank Dr. Congmao Wang for ongoing discussions. This study was supported in part by The Chinese National Transgenic Plant Special Funds (2016ZX08012-005), and the Program for Professor of Special Appointment (Eastern Scholar) at Shanghai Institutions of Higher Learning.

## Conflict of interest

There are no conflicts to declare.

## Data availability

The raw sequence data have been deposited in NCBI BioProject under accession code PRJNA757590.

## Supplemental File

### Supplemental Tables

**Supplemental Table S1.** The detail components of the seven tested samples.

| Sample name | GM event | Species | GM event content<br>(%, mass/mass ratio) |
| --- | --- | --- | --- |
| S1 | NK603 | maize | 50% |
| S2 | Mon810 | maize | 50% |
| S3 | TT51-1 | rice | 100% |
| S4 | GTS 40-3-2 | soybean | 5% |
| S5 | RT73, RF2 | canola | RT73 (99%), RF2(1%) |
| S6 | Non-GM | non-GM maize,<br>rice, soybean,<br>and canola | / |
| S7 | TT51-1 | rice | 100% |

**Supplemental Table S2.**
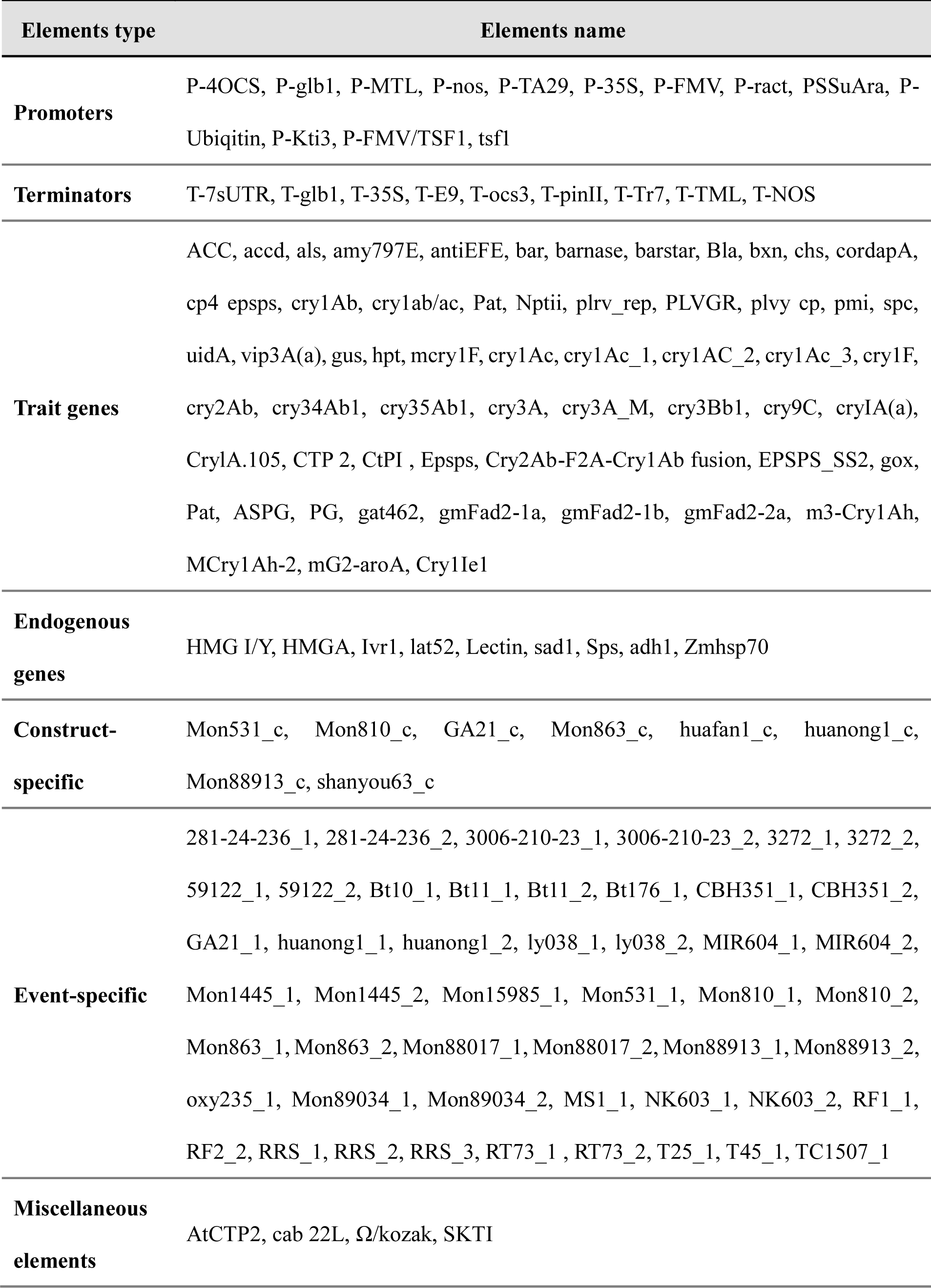
The elements used for the design of universal tiling probe panel.

**Supplemental Table S3.** Summary of designed universal tilling probe panel.

| Statistics | Number | Length (bp) | Probe Coverage (%) | Estimated Coverage (%) |
| --- | --- | --- | --- | --- |
| Designed targets | 150 | / | / | / |
| Universal probe panel Covered targets | 150 | / | / | / |
| Designed targets bases | / | 175038 | / | / |
| Universal probe panel target bases covered | / | 171678 | / | / |
| Universal probe panel Coverage | / | / | 98.08 | 99.64 |

**Supplemental Fig. S1.**
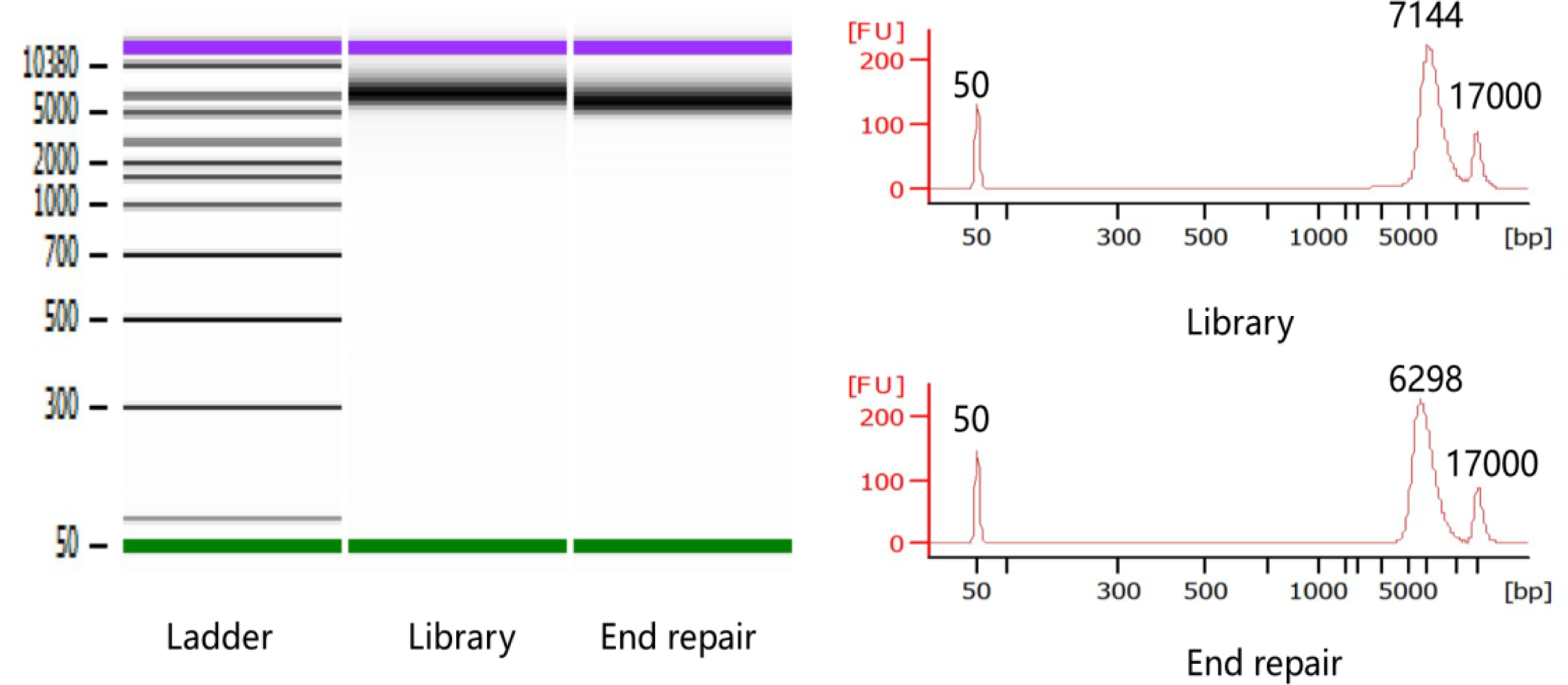
The quality evaluation of final constructed sequencing.

**Supplemental Fig. S2.**
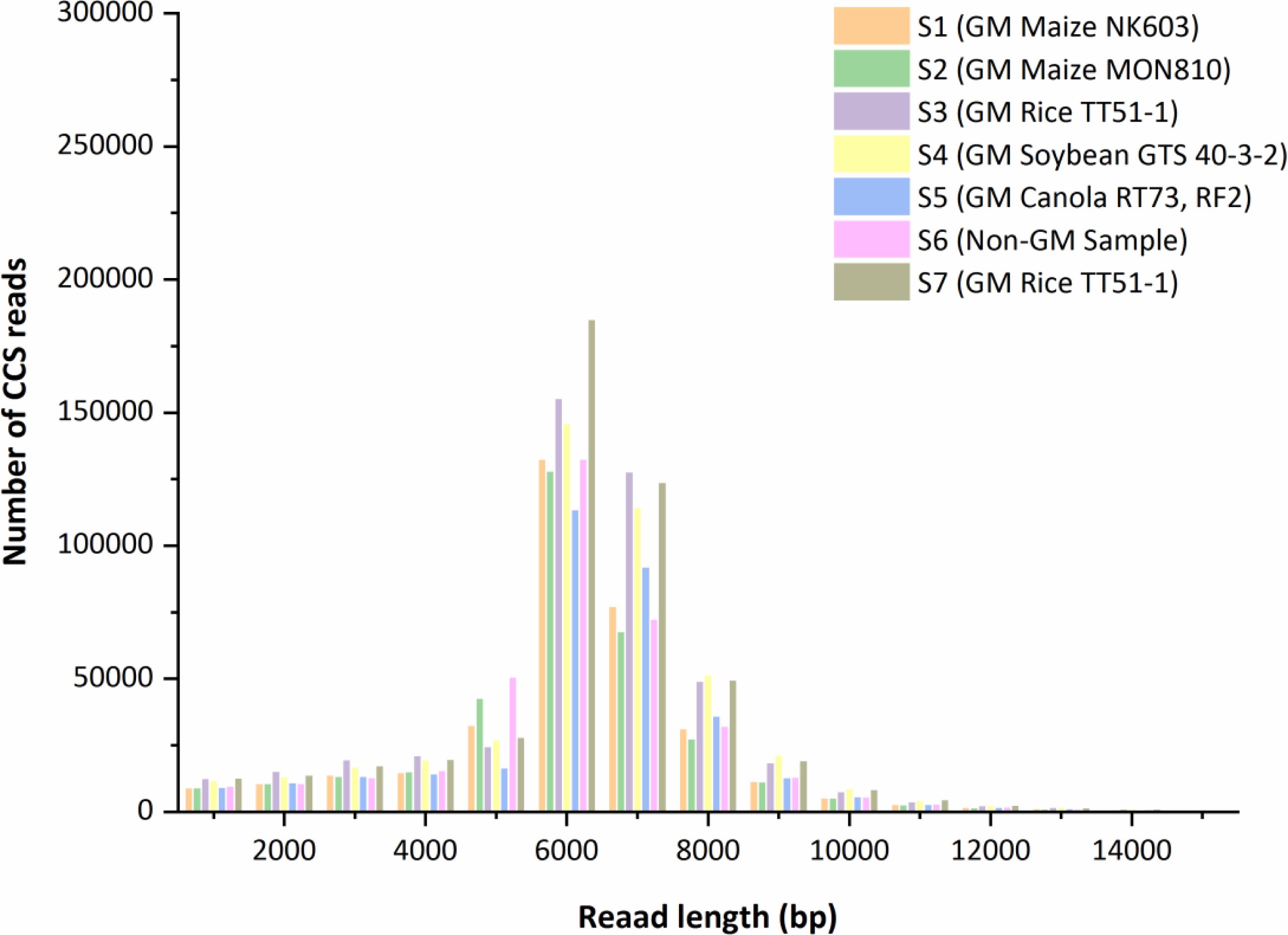
The length and number distribution of sequenced CCS reads of tested samples.

**Supplementary Figure S3.**
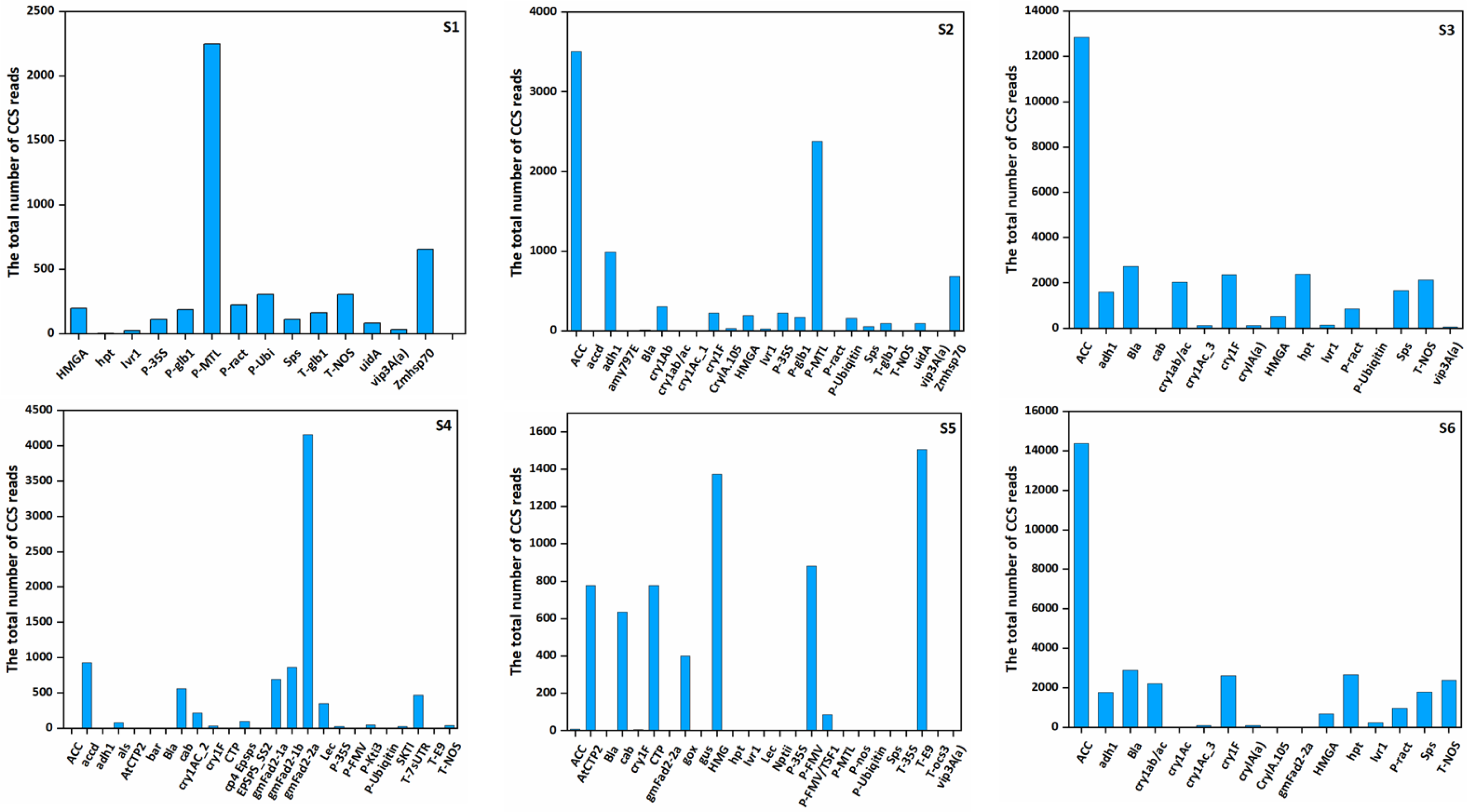
Statistical analysis of CCS reads mapped to transgenes and endogenous reference genes.

**Supplemental Fig. S4.**
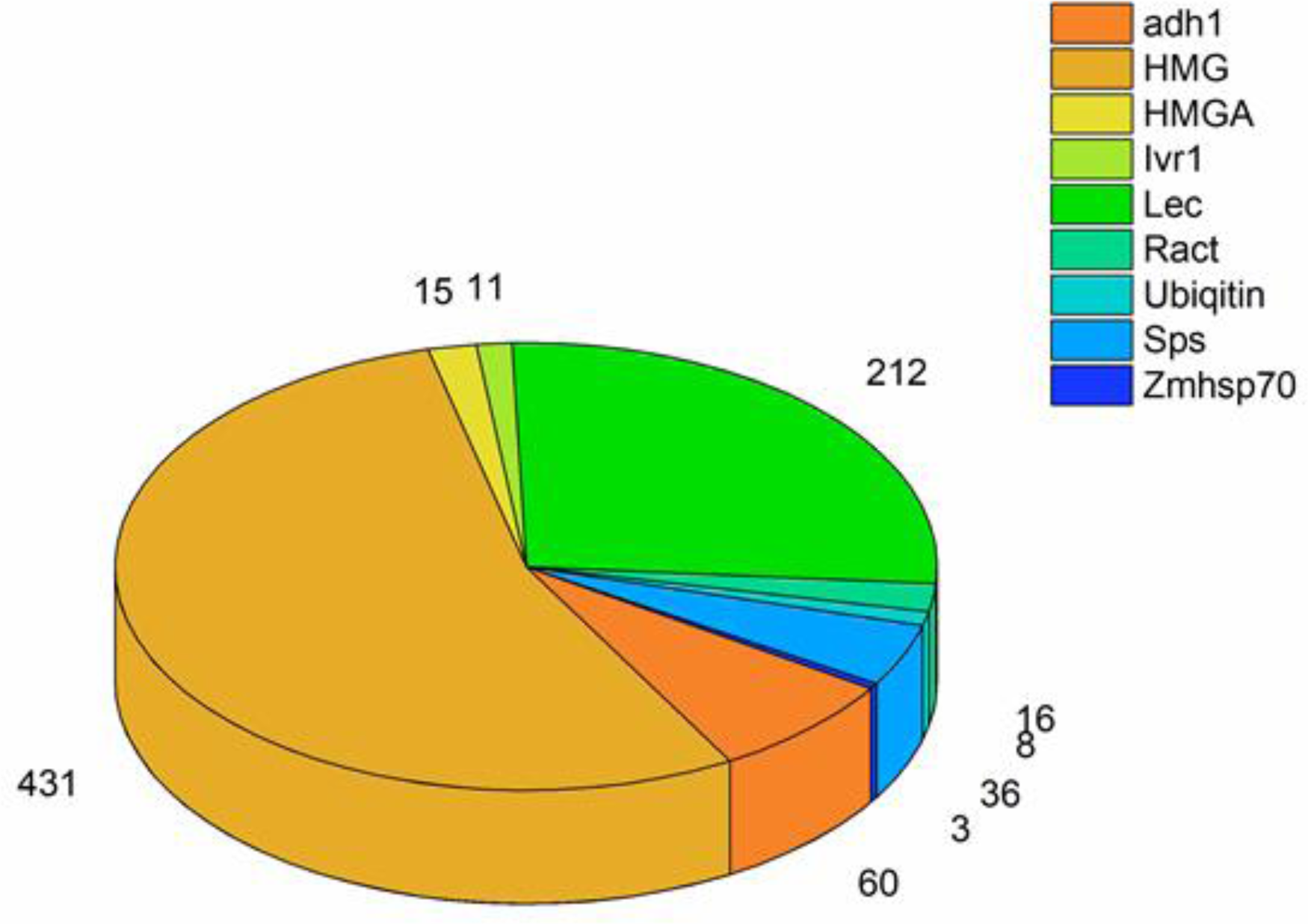
The distribution of the CCS reads which aligned to varied host native DNAs in sample S6.

## Notes

### Competing Interest Statement

The authors have declared no competing interest.

